# Extraction of individual associative memory related dominant theta frequency for personalized transcranial brain stimulation

**DOI:** 10.1101/2022.03.07.483124

**Authors:** Jovana Bjekić, Dunja Paunovic, Marko Živanović, Marija Stanković, Inga Griškova-Bulanova, Saša R. Filipović

**Affiliations:** Human Neuroscience Group, Institute for Medical Research, University of Belgrade, Serbia; Institute of Psychology and Laboratory for Research of Individual Differences, Department of Psychology, Faculty of Philosophy, University of Belgrade, Serbia; Institute of Biosciences, Life Sciences Centre, Vilnius University, Lithuania

## Abstract

Non-invasive brain stimulation (NIBS) has gained increased interest in research and therapy of associative memory (AM) and its impairments. However, the one-size-fits-all approach yields inconsistent findings, thus putting forward the need for the development of personalized frequency-modulated NIBS protocols to increase the focality and the effectiveness of the interventions. There have been only a few attempts to deliver theta frequency-personalized tES. The current study explores the feasibility of determining dominant individual theta-band frequency (ITF) based on AM task evoked EEG activity. In a sample of 42 healthy young adults, we extracted the frequencies (2-15 Hz, in 0.5 Hz steps) with the highest event-related spectral perturbation from the EEG recorded during successful encoding in the AM task. The developed method for extraction of the dominant theta-band frequency based on the AM-evoked EEG changes is able to reliably determine the AM-related ITF and can be used for personalization of the oscillatory NIBS techniques.

## Background

Brain oscillations arise from synchronized activity of large neuronal populations and have been associated with a variety of cognitive functions (Başar et al., 2001). The functional significance of rhythmic brain activity led to the increased interest in using non-invasive brain stimulation (NIBS) techniques which allow for direct modulation of brain oscillations (Herrmann et al., 2016). Specifically, transcranial electric stimulation (tES) techniques, such as transcranial alternating current stimulation (tACS) and oscillatory transcranial direct current stimulation (otDCS), have received considerable attention (Antal & Paulus, 2013; Fröhlich et al., 2015; Herrmann et al., 2013). In both of these techniques, weak sinusoidally modulated electric current is delivered to affect endogenous neural oscillations (Antal & Herrmann, 2016; Bland & Sale, 2019) and consequently, affect performance via network-wide recruitment (Tavakoli & Yun, 2017). The tACS/otDCS can affect brain activity either by matching the exogenous with intrinsic frequencies to increase the amplitude of the intrinsic oscillations (resonance), or by constraining/entraining brain oscillatory activity into desired frequency by stimulating regardless of the frequency of the intrinsic oscillations (Antal & Herrmann, 2016; Klink, Paßmann, et al., 2020).

Oscillatory brain activity in different frequency bands (i.e., delta (0.5–4Hz), theta (4–8Hz), alpha (8–12Hz), beta (12–30Hz), and gamma (30–80Hz)) have been related to the different cognitive processes. The majority of tACS/otDCS studies used mid-band frequencies across all participants e.g., 10Hz for “alpha-band stimulation” (for review see (Klink, Paßmann, et al., 2020; Schutter & Wischnewski, 2016) to modulate cognitive functions. However, this *one-size-fits-all* approach yielded inconsistent findings and put forward the need for personalized frequency-specific NIBS in research and therapy (e.g., see(Figee & Mayberg, 2021; Frohlich & Riddle, 2021). This entails tuning the stimulation waveform to the endogenous network dynamics by using the individual peak frequency of the targeted oscillation (Frohlich & Riddle, 2021). Here we focus on the challenge of personalizing transcranial electrical stimulation (tES) for associative memory (AM) enhancement using AM task-related EEG data to determine person-specific dominant theta-band frequency. The focus is put on AM – the ability to remember multiple pieces of information, binding them together, and encoding them as a meaningful unit. The AM impairment is one of the most prominent early symptoms of dementia and mild cognitive impairment (Bastin et al., 2014; Delhaye et al., 2019) and thus an important target-function for the development of personalized NIBS based treatments.

Theta oscillations dominate hippocampal electrical activity (Nuñez & Buño, 2021) and are considered to play a critical role in the hippocampal-neocortical interactions (Hanslmayr et al., 2016; Sirota et al., 2008) Plochl et al 2020 as well as in the interactions across the widespread neocortical circuits (Zhang et al., 2018). Theta oscillations have long been implicated in learning and memory (Herweg et al., 2020). Namely, theta activity was found to be associated with processes of information and contextual processing (Kragel et al., 2020), temporal organization for memory engrams (Lisman & Jensen, 2013; Turi et al., 2018), and associative binding (Clouter et al., 2017). Furthermore, theta activity has been observed during lower-level mnemonic processes such as encoding, recognition, and recall (Beppi et al., 2021). Recent reviews argue that theta-band oscillations are in fact causally engaged in AM (Herweg et al., 2020).

Building up on that, previous NIBS studies aiming at memory neuromodulation used theta-band frequencies to entrain cortico-hippocampal circuits. Most of these studies used a single stimulation-frequency within theta band across all participants: either 4Hz (Alekseichuk et al., 2020; Bender et al., 2019), 5Hz (Kleinert et al., 2017; Klink, Peter, et al., 2020; Vulić et al., 2021) or 6Hz (Abellaneda-Pérez et al., 2020; Alekseichuk et al., 2017; Antonenko et al., 2016; Lang et al., 2019; Lara et al., 2018; Polanía et al., 2012; Röhner et al., 2018; Tseng et al., 2018; Violante et al., 2017). However, the evidence suggests that the choice of stimulation frequency may be relevant factor that can modulate the NIBS effects. Namely, two studies which contrasted effects of low (4Hz) and high (7Hz) theta tACS found differential effects on memory (X. Guo et al., 2021; Wolinski et al., 2018). One of the reasons for these differential effects may lay in the individual differences in participants’ peak or dominant theta rhythm – it has been suggested that the stimulation effects may be the most prominent when the stimulation frequency is close to a person’s endogenous peak or dominant frequency (Kleinert et al., 2017; Stecher & Herrmann, 2018).

There have been only a few attempts to deliver theta frequency-personalized tES (Jaušovec et al., 2014; Pahor & Jaušovec, 2014, 2018; van Driel et al., 2015). They used different methods to determine endogenous peak theta frequency or, so-called, individual theta frequency (ITF). The first method relied on the most pronounced peak in the power spectrum of the resting state EEG – the peak of the posterior background activity, i.e., alpha peak, or, so called, “individual alpha frequency” (IAF). The IAF is then used as an anchor for determining personalized windows for other frequency bands (Klimesch, 1999). Following this approach, Jaušovec and colleagues (Jaušovec et al., 2014; Pahor & Jaušovec, 2014), applied oscillating currents in ITF for working memory neuromodulation. In their studies ITF was determined as resting state IAF-5Hz (e.g., person 1: IAF = 9.5 à ITF = 4.5; Person 2: IAF= 11 à ITF = 6, etc.). The second approach to determine ITF is based on cross-frequency theta-gamma coupling. It relies on the cross-frequency covariance and is built upon the evidence on theta-gamma phase coupling in the hippocampus as a direct substrate of memory processes (Lisman & Jensen, 2013; Sirota et al., 2008). Determining ITF based on theta-gamma frequency coupling assumes finding the theta band frequency with the highest correlation with gamma band frequency, which is usually done through quantifying phase–amplitude coupling (Onslow et al., 2011). This approach has been adopted by different studies aiming to modulate working memory using NIBS (see (Abubaker et al., 2021) for review). Finally, the most straightforward approach to extract ITF would be to find the theta band frequency with the highest power during function-relevant task. This approach was adopted by van Driel and colleagues (van Driel et al., 2015) who used automatic peak frequency detection method to extract theta frequency with the maximal power (log transformed and detrended to attenuate 1/f power scaling) from the EEG activity during the cognitive inhibition task.

However, none of the previous studies delivered ITF-personalized NIBS to enhance AM, and there is no evidence that any of these ITF extraction approaches captures AM-relevant EEG activity. Furthermore, the ITF extraction methods have not been described in sufficient detail, and often not critically evaluated. For example, reporting the number of participants for which ITF could not be reliably extracted and the distribution of extracted ITFs are usually omitted. Finally, technical challenges of frequency-personalized NIBS are rarely addressed, hence the solutions to enable reproducibility across different labs are largely unavailable.

To set the ground for future basic research and clinical trials, the current study explores the feasibility of ITF-personalizing. Specifically, we explore the feasibility of determining dominant individual theta-band frequency based on AM task evoked EEG activity.

## Method

### Study design

The study reported here was carried out as a part of a large project aimed at assessment of the several personalized oscillatory tES techniques on associative memory. The study was in fact a session zero, used to determine the individual associative memory related theta frequency over parietal regions, and was followed by a series of interventional sessions in each of them a different tES technique or sham were applied. (Figure 1).

**Figure 1.**
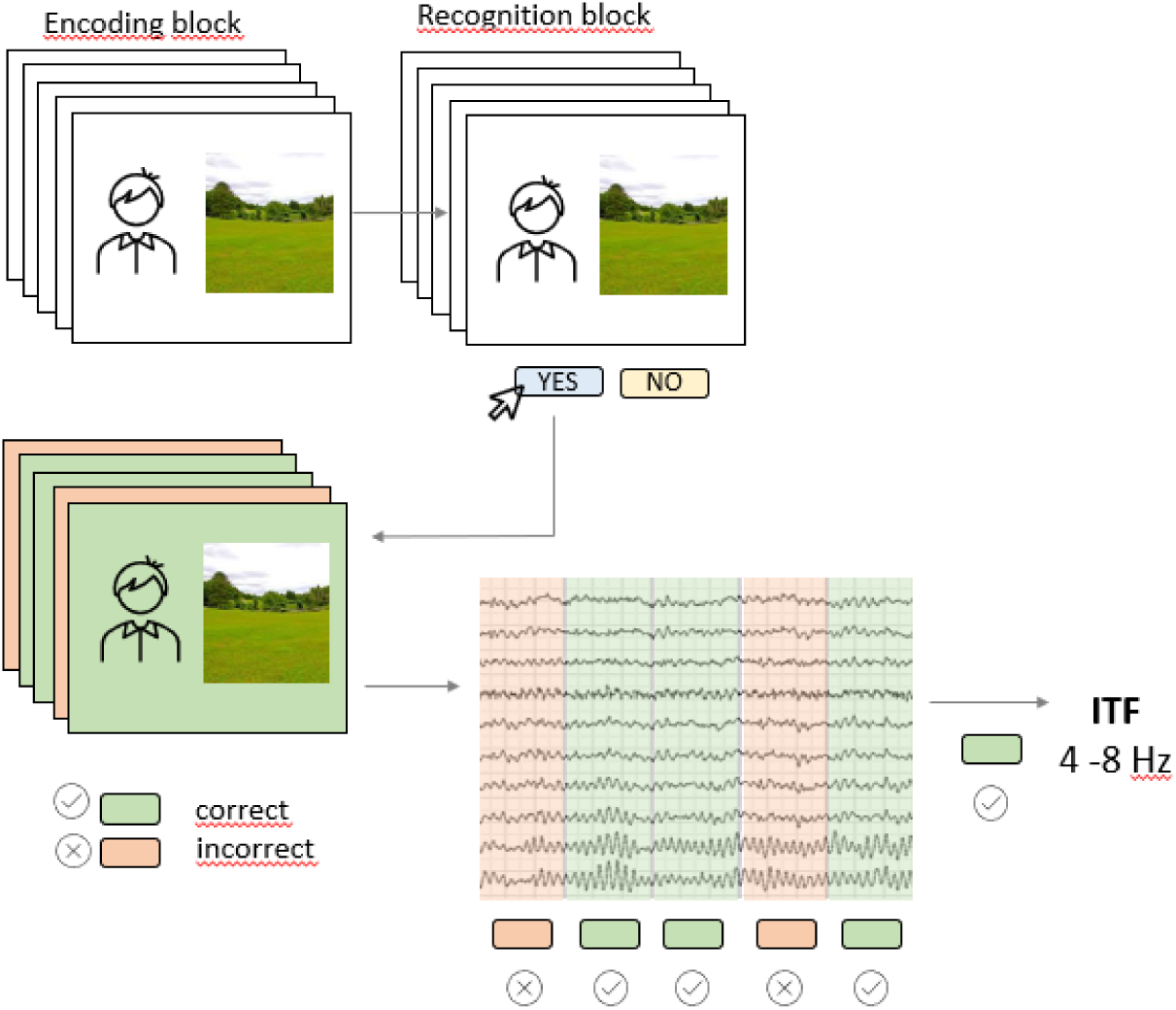
The AM task (encoding and recognition block) was performed while EEG was recorded; the EEG signal form the encoding bock was used to extract individual theta frequency for subsequently correctly recognized pairs.

Before the inclusions, all volunteers were screened by an online assessment questionnaire for the inclusion and exclusion criteria. Once the criteria were met, the volunteers were contacted and the first recording session (i.e., the session zero) was booked. The participants were instructed to abstain from alcohol for 24h, and from nicotine and caffeine-containing dinks at least for one hour before the session.

At the recording session, after signing the informed consent, the participants were first familiarized with the setting and then had the EEG recording during part of which they completed the AM task. The EEG recordings obtained during the AM task were subsequently used to extract ITF.

### Participants

Forty-two young adults (age: 22 – 34 years, *M* = 25.05, *SD* = 3.55; 26 female) took part in the study. All participants were right-handed (Edinburgh Handedness Inventory (Veale, 2013) laterality quotient > 80), with normal or corrected-to normal vision and they all satisfied common tDCS inclusion/exclusion criteria – i.e., reported no history of neurological or psychiatric disorders, traumatic brain injury, metal implants in the head. All participants gave their written informed consent and were compensated for their involvement in the study. The study was conducted in line with the guidelines of the Declaration of Helsinki and was approved by the Institutional Ethics Board (EO129/2020).

### Associative memory task

To assess AM, we used visual paired associates task consisting of encoding and associative recognition block (Figure 1). In the Encoding block, 42 face-scene pairs were presented successively for 2000ms, and participants were instructed to remember them as pairs. The inter-stimulus-interval (ISI) was randomly varying between 1250ms and 1750ms, and during this time a white screen with a black fixation dot was presented. The stimuli were portrait pictures of young Caucasian’s of both sexes taken from FEI database (Thomaz & Giraldi, 2010) while scenes were publicly available pictures of all-natural scenery such as forests, seacoasts and fields. In Recognition block, participants saw 84 face-scene pairs, half of which were correctly paired i.e., the same pairs they have previously seen in Encoding block, while the other half were recombined pairs that consisted of wrongly paired faces and scenes presented in the Encoding block. The pairs were presented successively, and participants’ task was to recognize the pair as either “old” or “recombined” by pressing one of the assigned keyboard keys. The full task code with integrated EEG triggers is available at https://osf.io/be8df/. The AM task was programmed and administered in OpenSesame software (Mathôt et al., 2012) and presented on a 23-inch monitor (0.6m head-to-screen distance).

### EEG recording

To record EEG, we used light, mobile, battery-operated, hybrid tDCS-EEG Starstim device (Neuroelectrics Inc., Barcelona, Spain), which was remotely operated via Neuroelectrics® Instrument Controller (NIC2) software (Neuroelectrics Inc., Barcelona, Spain).

EEG was recorded with Ag/AgCl electrodes (4 mm diameter, 1 cm^2^ gel-contact area) from 20 positions (Fp1, Fp2, Fz, F3, F4, F7, F8, T7, T8, Cz C3, C4, CP5, CP6, Pz, P3, P4, PO7, PO8, and Oz according to the international 10-10 EEG positioning system). For the reference (CMS) and ground (DLR), we used either ear-clip with dual CMS-DLR electrode on the right earlobe or pre-gelled adhesive electrodes on the right mastoid, depending on the signal quality. The impedance was kept below 5kΩ throughout the recording. The EEG signals were recorded with the sampling rate of 500Hz, 0 – 125Hz (DC coupled) bandwidth, and 24 bits – 0.05μV resolution.

The offline EEG preprocessing was performed in EEGLAB for MATLAB (Delorme & Makeig, 2004). The signal was high-pass filtered at 0.1 Hz and the power-line noise (50Hz) was removed using multi-tapering and Thomas F statistics, as implemented in CleanLine plugin for EEGLAB. The channels with substantial noise were excluded after eye inspection. The independent component analysis (ICA), using “runica” routine with default settings, was performed on the remaining channels to detect and remove eye movement artefacts.

The EEG signal recorded during the encoding block of the AM task was analyzed. The epochs were created from -1000ms to 2500ms in respect to the stimulus onset, and baseline-corrected to the pre-stimulus period (−800ms to -100ms). The data were visually inspected, and bad epochs were manually rejected. The epochs were labeled based on the subsequent recognition accuracy – namely, the epochs containing encoding stimuli that were correctly identified as “old” in the recognition block were labeled as “successful encoding” and the trials that were incorrectly identified as “recombined” were labeled as “unsuccessful encoding”.

### Individual theta frequency

Since in this study we were interested in the AM related theta activity, the ITF was defined as the dominant theta-band frequency during successful AM encoding. Therefore, the ITF extraction was performed on “successful encoding” epochs, within the time-window 250ms to 1250ms from the stimulus onset, since this is the time-window when AM processes related EEG activity has been usually recorded (Friedman & Johnson, 2000).

Signal was baseline-corrected to the mean of the pre-stimulus period. It was further subjected to complex Morlet wavelet (7 cycles) to extract frequencies from 1 to 15 Hz in 0.5 Hz resolution using MATLAB Wavelet Toolbox. To calculate the event-related spectral perturbation (ERSP), indicating event-related changes in power relative to a pre-stimulus baseline, we used the formula:

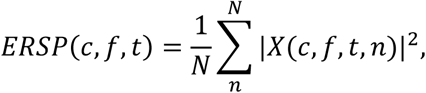

where for every channel c, frequency f, and time point t a measure is calculated by taking time frequency decomposition X of each trial n (Mørup et al., 2007). The ERSP was further expressed as a ratio against the baseline (−800 ms to -100 ms) for each 0.5 Hz, starting from 2 to 15Hz. Next, using custom written MATLAB script, the frequency with the highest ERSP value was extracted from each of 19 overlapping time windows (100ms width in 50ms steps; from 250ms to 1250ms; i.e., 250ms-350ms, 300ms-400ms, 350ms-450ms, … 1150ms-1250ms) at each of six centroparietal electrodes (Cz, C3, C4, Pz, P3, P4). The selected electrodes cover the scalp area where the AM related EEG activity is known to be expressed the best (Guo et al., 2005). Finally, to extract dominant theta band frequency, we calculated the mode (i.e., the most frequently occurring value) for the frequencies between 4-8Hz (in 0.5Hz steps) in the time x electrode matrix for each participant (114 cells per participant).

### Statistical analysis

The statistical analyses were performed using Analyze Data module in Microsoft Excel and in JASP. As the measure of AM performance, we calculated the number and the percentage of correctly identified targets and correctly rejected recombined pairs as well as the overall success rate. The ITF was extracted as modal frequency from electrode x matrix of each participant (see 2.4.3). Descriptive statistics, such as mean, standard deviation, median, range etc. were calculated for all variables. In addition to that, the relative share of theta-band frequencies in electrode x matrix were calculated as well as the relative share of the ITF in cells with theta-band frequencies. The later was interpreted as the participant-level reliability of the extracted ITF.

## Results

The overall success rate in AM task was on average 64.0% (*SD* = 7.70), with average of 68.5% (*SD* = 13.19) for correctly rejected recombined pairs and 59.5% (*SD* = 12.72) for correctly identified targets (Figure 2a). As encoding epochs were selected based on subsequent associative recognition (“successfully encoded”), this resulted in 25 epochs on average (out of 42 in total) to be used for ITF extraction. However, due to substantial variability in memory performance the number of epochs on the individual level was between 15 and 37. After removal of bad epochs from EEG signal, between 14 and 37 *(M* = 23.57, *SD* = 5.71) epochs remained for the analysis.

**Figure 2.**
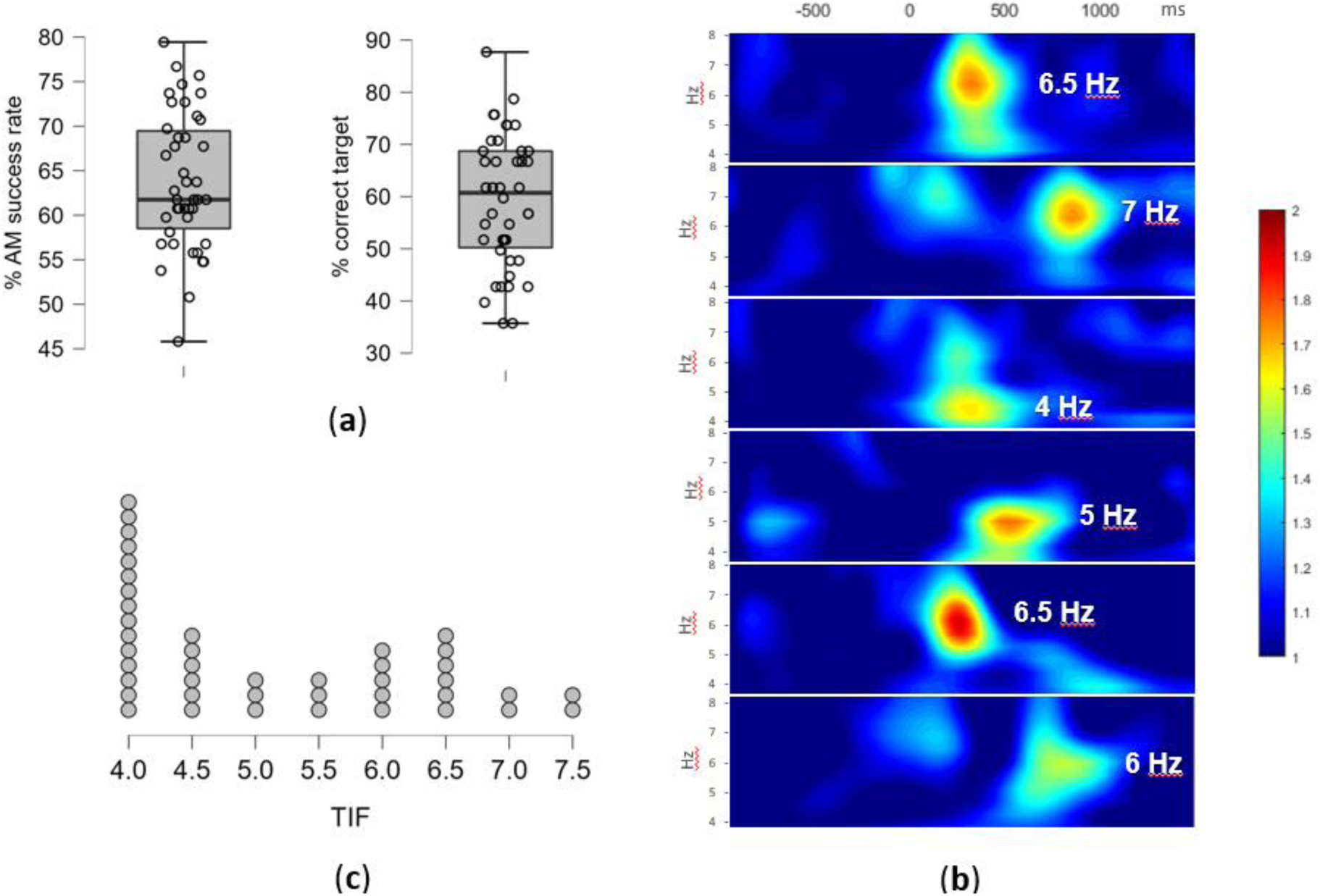
**(a)** The AM task performance – the overall success rate in % and the % of correctly identified targets i.e., successfully encoded pairs. Individual participants overlay the box-plot marking median and interquartile range (**b**) The individual differences in theta activity during AM encoding - examples of time-frequency analysis of EEG signal during the successfully encoded pairs for six participants; the ERSP values, averaged across 9 centro-parietal electrodes, are plotted for the time-window between -800 and 1250ms (x-axis) and for theta-band frequencies 4-8Hz (y-axis), with the extracted ITF for each participant marked; (**c**) The individual theta frequencies (ITF) distribution – the number of participants and their extracted ITF values in Hz (x-axis).

The individual differences in peak theta-frequency as well as the time-range of enhanced theta- band activity occurrence during successful AM encoding are presented in selected time *x* frequency plots on Figure 2b. The time *x* electrode matrix for each participant showed high inter-individual variability in theta-activity dominance i.e., the number of cells with highest ERSP in theta in comparison to neighboring frequency-bands i.e., delta and alpha. The ERSP peaks in theta-band frequencies have been observed in 51.5% of cell on average (*SD* = 24.1%), with the high inter-individual variability (range: 7.9% - 92.1%)

The ITF, defined as modal theta-band frequency, was successfully extracted for 39 participants (93%), while for the remaining three, additional steps in the analyses had to be performed. Namely, one participant had double mode distribution of frequencies (7.5Hz between 650-950ms and 5.5 Hz between 950-1200ms on all electrodes), and two participants did not have a prominent theta activity in the defined time *x* electrode matrix, thus EEG signal from additional electrodes (P7 and P8) was analyzed to extract ITF. Eventually, ITF ranged between 4Hz and 7.5Hz (*M* = 5.16, *SD* = 1.16), with the distribution as presented in Figure 2c.

However, it is important to note that there was significant variability between participants in homogeneity of peak theta-frequencies i.e., in occurrence of ITF within theta-band frequencies during successful AM encoding. Namely, the ITF was the peak theta-frequency in on average 0.53 (*SD* = 0.20; range: 0.26 – 1.00) of theta-band peaks. Based on the incidence of their ITF, the participants (and therefore extracted ITFs) could be split into the following groups: singular ITF [> 0.80 (*n* = 5)], highly reliable ITF [0.80 -0.51 (*n* = 13)], reliable ITF [0.50-0.31 (*n* = 18)], unreliable ITF [0.30-0.15 (*n* = 8)], and no ITF [< 0.15. i.e., below the chance level (*n* = 0)].

## Discussion

The study shows the feasibility of extracting ITF that reflects AM-relevant neurophysiological activity. To the best of our knowledge, this is the first attempt to extract ITF from AM-task related EEG.

We conceptualized ITF as the dominant theta-band frequency during successful AM encoding. Hence, to capture individual differences in AM encoding time-frequency spectra, the new approach for ITF extraction needed to be developed. Similar to van Driel and colleagues (van Driel et al., 2015), we opted for an approach that is based on the differences in power between theta-band frequencies. However, since automatic peak detection cannot ensure that identified ITF has sufficiently higher power than other frequencies, and since the latencies of theta synchronization differed between participants, we decided to extract peak frequencies across multiple time windows and multiple electrodes. This does not change the basic assumptions of power-based approach but provides a more reliable ITF estimate due to the increased number of observations. The ITF extraction based on the modal theta-band frequency across multiple time-windows and electrodes resulted in 93% of successfully identified dominant frequencies, while the remaining cases could be resolved by additional analysis of neighboring electrodes. Still, it is important to note that even with the relatively large number of observations (more than 100), it can be difficult to identify a single modal value. This opens the question if the ITF-based personalization of the tACS/otDCS should be understood as a continuum, that is, if we need to assess and report the level of NIBS personalization based on the reliability of ITF. This ITF extraction approach allows for that possibility by simple calculation of the proportion of cells in time *x* electrode matrix in which ITF value appears for each participant. Namely, if we treat value (i.e., frequency with the highest ERSP) within the time *x* electrode matrix as a repeated measure of the same underlying process responsible for successful encoding, the proportion of occurrence of each value can be interpreted as their reliability in repeated measurement. For example, if an ITF occurs in .70 of the “measurements” across time and electrodes, a given value can be considered as highly reliable and therefore inherent for a successful encoding of a given individual. Consequently, when used as an input parameter for tACS/otDCS, the protocol can be considered as highly personalized. Conversely, if a participant’s ITF occurs in fewer than 1/3 of the total measurements, i.e., has the reliability of < .30, a given value cannot be regarded as reliable and thus ITF based tACS/otDCS protocol could not be labeled as well- personalized.

Since ITF was extracted from task-related EEG signal, several aspects of the AM task itself need to be put forward. First, the AM was operationalized by paired-associates task typically used in AM NIBS studies (Bjekić, Čolić, et al., 2019; Bjekić, Vulić, et al., 2019; Flöel et al., 2012; Leach et al., 2019; Leshikar et al., 2017; Matzen et al., 2015; Zhang et al., 2018). The task was designed to be of the optimal difficulty for young healthy volunteers, and as such it cannot be used in older adults or clinical populations without adjustments or prior testing. Furthermore, the individual differences in AM performance resulted in variable number of epochs to be analyzed. The number of epochs may be smaller than what is recommended for reliable EEG analysis. However, this issue cannot be addressed simply by increasing the number of encoding trials as that would increase the difficulty of the task and probably result in higher chance-based responses in the recognition phase. Second, we opted for analyzing the EEG signal from encoding rather than recognition block, based on the assumption that theta-rhythm underlies the process of binding – that is, creating new associations in memory (Kota et al., 2020; Lega et al., 2012; Lin et al., 2017), and that the encoding EEG is free of other processes involved in the task-based- recognition (e.g., decision making, motor-control/action). Finally, the stimuli were selected to be as ecologically valid as possible (human faces and landscapes), however, it cannot be claimed that the same ITFs would be extracted from same-structure task using different type of stimuli.

Despite these constrains, it could be argued that the proposed ITF extraction method is superior to the alternative approaches available in the literature such as analyzing individual differences in resting state EEG (Jaušovec et al., 2014) and theta-gamma cross-frequency coupling ITF (Pahor & Jaušovec, 2018), at least when comes to NIBS for AM. Namely, even though ITF defined as IAF-5Hz ensures that the most prominent peak is used as anchor (Klimesch, 1999), it assumes universal equidistance between IAF and ITF, which may not be the case. More importantly, it could be argued that ITF for memory should not be directly extrapolated from resting state EEG due to the individual differences in neural oscillators engaged at rest or during task performance. The cross-frequency coupling method has stronger theoretical background as there is evidence that theta-gamma coupling in the hippocampus underlies memory processes (Lisman & Jensen, 2013; Sirota et al., 2008). However, this theta-gamma coupling based ITF has not been evaluated in AM studies. Moreover, phase– amplitude coupling is only one of the many possible implementations of cross-frequency coupling in neuromodulation. Namely, from the theoretical perspective in addition to phase-amplitude coupling the cross-frequency coordination can also be power-to power; phase-to-phase (or phase-locking), phase-to-frequency (Jensen & Colgin, 2007). As the recent review concludes that the relationship between different coupling phenomena and memory are still understudied (Abubaker et al., 2021) this approach currently seems promising for the development of different long-range network multichannel tES protocols and assessing their effects (see e.g., (Alekseichuk et al., 2016), but it should be used with caution for extracting ITF to serve as input parameter for personalizing NIBS protocol. Finally, none of the previously used ITF extraction methods allowed for quantifying the participant-level reliability of the ITF and consequently the level of personalization achieved by frequency-tuning tACS/otDCS.

## Conclusion

The developed method for extraction of dominant theta-band frequency based on AM task evoked EEG changes can be used to reliably determine the AM-task-related ITF that can be used for personalization of oscillatory NIBS techniques.

